# AGES for Ageing: Evaluating the auxin-inducible gene expression system for use in *Drosophila* ageing studies

**DOI:** 10.64898/2026.05.28.728476

**Authors:** Taylor McGilvary, Kriti Gupta, Adam J. Dobson, Nathan Woodling

## Abstract

Our research is only as good as our tools allow it to be. The fruit fly *Drosophila* has been a fundamental discovery platform in uncovering evolutionarily conserved biological underpinnings of ageing, due in large part to an ever-expanding functional genetic toolbox which permits fine-tuneable and cell-type-specific modulation of gene expression with relative ease. However, many existing gene expression systems present limitations for studying fly ageing, including off-target effects for inducing agents that allow temporal control. More recently-generated tools such as the auxin-based gene expression system (AGES) therefore present opportunities as potential alternatives in our methodological repertoire for ageing research. Here we have evaluated the AGES system in a variety of contexts in *Drosophila* ageing. We find that AGES can effectively induce transgene expression across a range of ages, albeit with tissue-specific efficiency. However, we also observe several phenotypes from auxin feeding, even in non-AGES genotypes, that may confound studies focused on ageing research, including reduced body mass and reduced survival under starvation and oxidative stress conditions. We also observe phenotypes from activating the AGES machinery, including shortened lifespan, that could present challenges for using AGES in longevity-based studies. Nevertheless, we find that AGES can be used to recapitulate at least some effects of well-established pro-longevity interventions – for instance reduced fecundity from expression of a dominant-negative form of the insulin receptor – reinforcing the value of AGES in certain domains. Taken together, our results underscore the need for caution and comprehensive controls in ageing studies that rely on functional genetics, regardless of the chosen genetic toolset.

## Introduction

The fruit fly *Drosophila melanogaster* has been indispensable over the past century in research to unravel the biological principles of ageing. This includes our understanding of ageing as underpinned by specific evolutionarily conserved signalling pathways. For instance, *Drosophila* research has revealed that healthy lifespan can be extended through multiple interventions, including perturbation of insulin and/or TOR signalling (Clancy *et al*., 2001; Tatar *et al*., 2001; Kapahi *et al*., 2004; Proshkina *et al*., 2015), restriction of specific dietary components (Partridge, Piper and Mair, 2005; Kapahi, Kaeberlein and Hansen, 2017), and tissue-specific interventions such as enhanced neuronal autophagy (Simonsen *et al*., 2008; Ulgherait *et al*., 2014; Bolukbasi *et al*., 2021) or altered intestinal stem cell proliferation (Biteau, Hochmuth and Jasper, 2008; Rera, Clark and Walker, 2012).

These advances, in large part, have been made possible by an ever-expanding genetic toolbox in *Drosophila*, which permits fine-tuneable, tissue- and cell-type-specific modulation of gene expression, and thus biological function, with relative ease. This toolkit has largely been built upon the backbone of multi-partite expression systems such as the classical GAL4/GAL80/UAS system, which utilises a transcriptional activator (*GAL4*) and/or its repressor *GAL80* placed under the control of specific promoters to mimic the spatial and temporal expression pattern of those promoters when combined with a complementary responder gene (*UAS-transgene*) (Brand and Perrimon, 1993). Further iterations of this system, namely the temperature-dependent GAL80^ts^ system (TARGET) (McGuire *et al*., 2003) and the drug-inducible GeneSwitch system (Osterwalder *et al*., 2001), have since added conditional control over transgene expression, for example allowing researchers to bypass developmental effects of genetic perturbations while permitting investigation of their role in ageing via adult-onset manipulation (Sun *et al*., 2013). However, current tools present some limitations for studying ageing in the fly, due to off-target effects of inducers used. For instance, given the poikilothermic nature of *Drosophila*, temperature can affect metabolic rates and longevity, which can render use of the TARGET system challenging due to possible confounding effects (Miquel *et al*., 1976; Mołoń *et al*., 2020). Regarding the GeneSwitch system (He and Jasper, 2014), drawbacks include safety concerns around the antiprogestogen effects of the inducer compound RU-486, “leaky” transgene expression in the absence of RU-486 (Scialo *et al*., 2016), variability in expression levels achieved due to dose and age differences (Poirier *et al*., 2008), and issues regarding the palatability of RU-486 influencing feeding and lifespan in a diet-dependant manner (Yamada *et al*., 2017). Thus, these systems require careful selection of experimental controls for accurate interpretation of results.

A recently-developed system for conditional GAL4 expression is the auxin-based gene expression system (hereafter referred to as AGES), which relies on spatiotemporal modulation of gene expression via synthetic variants of the phytohormone auxin, typically via K-NAA (1-naphthaleneacetic acid potassium salt) (McClure *et al*., 2022). In brief, this system utilises GAL80 flanked by fused auxin-inducible degron (AID) tags, which, in the absence of auxin, accumulates in cells leading to inhibition of GAL4 transcriptional activity. After auxin ingestion, GAL80 is polyubiquitinated and targeted for degradation, alleviating GAL4 repression and permitting *UAS-transgene* expression. Crucially, major benefits include compatibility with vast existing libraries of *Drosophila GAL4* lines and auxin’s water solubility, cost effectiveness, and high safety profile.

Numerous studies have now successfully used the AGES system in fields including development (Corke *et al*., 2025), immune signalling (Kuklinski *et al*., 2026), and behaviour and metabolism (Richhariya *et al*., 2023, 2025; Fleck *et al*., 2024). However, the utility of AGES to study ageing has been less well explored, which we sought to redress in this study. Here we evaluate how transgene induction following auxin feeding varies by sex, chromosomal transgene location, auxin dose, and age. In addition, we explore whether AGES can recapitulate a classical anti-ageing intervention of reducing insulin signalling, examining both lifespan and healthspan parameters in this setting.

## Materials and Methods

### Fly stocks and husbandry

The white Dahomey strain of *Drosophila melanogaster,* positive for the bacterial cytoplasmic endosymbiont Wolbachia (*w^Dah+^)*, was the genetic background for all flies in this study. Wild-type Dahomey (*Dah*) flies were collected in 1970 in Dahomey (now Benin) and have since been maintained in large population cages with overlapping generations. The white Dahomey (*w^Dah^*) stock was derived by backcrossing the *w^1118^* mutation into this outbred background. Fly stocks used here were *da-GAL4* (FBti0013991) (Ikeya *et al*., 2009); *tubP-TIR1-2A-GAL80.AID* [‘AGES’] on the second chromosome (a kind gift of C. McClure, Queens University Belfast); *tubP-TIR1-2A-GAL80.AID* [‘AGES’] on the third chromosome (BDSC #92470, described in (McClure *et al*., 2022)); *UAS-InR^DN^* [*InR^K1405A^*] (BDSC #8252); and *UAS-mCD8::GFP* (BDSC #5130). *da-GAL4,UAS-mCD8::GFP* and *da-GAL4,tubP-TIR1-2A-GAL80.AID* [‘AGES’] stocks were recombined from their single constituents within the lab for this study.

All transgenes were backcrossed into the outbred *w^Dah^*^+^ background for six generations but have since been maintained as stocks and likely possess a degree of genetic background variability. For Figures 5 and 6 (lifespan and stress survival assays), however, parental stocks were refreshed into the *w^Dah^*^+^ background by performing backcrossing for an additional three generations immediately before experimental use.

Our protocol for experimental cultures is described in (Piper and Partridge, 2016). Briefly, all experimental flies were reared at a standardised larval density. Emerging adults were then collected within a 24-hour window and allowed to mate for 48 hours before being separated into single-sex vials at a density of 15 flies per vial, housed in DrosoFlippers (drosoflipper.com). Both stocks and experimental flies were maintained and experiments conducted at 25°C on a 12h:12h light:dark cycle.

### Fly media

All stocks and experimental flies were reared on sugar-yeast-agar (SYA) media containing 10% (w/v) brewer’s yeast (#903312, MP Bio), 5% (w/v) sucrose (#BP22010, Fisher Scientific) and 1.5% (w/v) agar (#A7002, Sigma) supplemented with 0.3% nipagin (w/v) (added as 30mL/L from a 10% (w/v) stock solution in ethanol (#H3647, Sigma), and 0.3% (v/v) propionic acid (#149300025, Fisher Scientific) as preservatives (Bass *et al*., 2007).

For auxin-based and corresponding control media, K-NAA (#GK2088, Glentham Life Sciences) and/or KCl (#26764.260, VWR Chemicals) were added from 2 M stock solutions in dH2O to standard SYA food, thoroughly mixed, and dispensed into individual fly vials to achieve the desired final concentrations as specified for each experiment. For experiments pertaining to Figures 1–3, auxin-based medium was supplemented with KCl, such that the final concentration of potassium across all conditions was 20mM. For experiments related to Figures 4–6, KCl was added to control media only, such that the resultant potassium concentration was 5mM in both control and auxin conditions.

**Figure 1.**
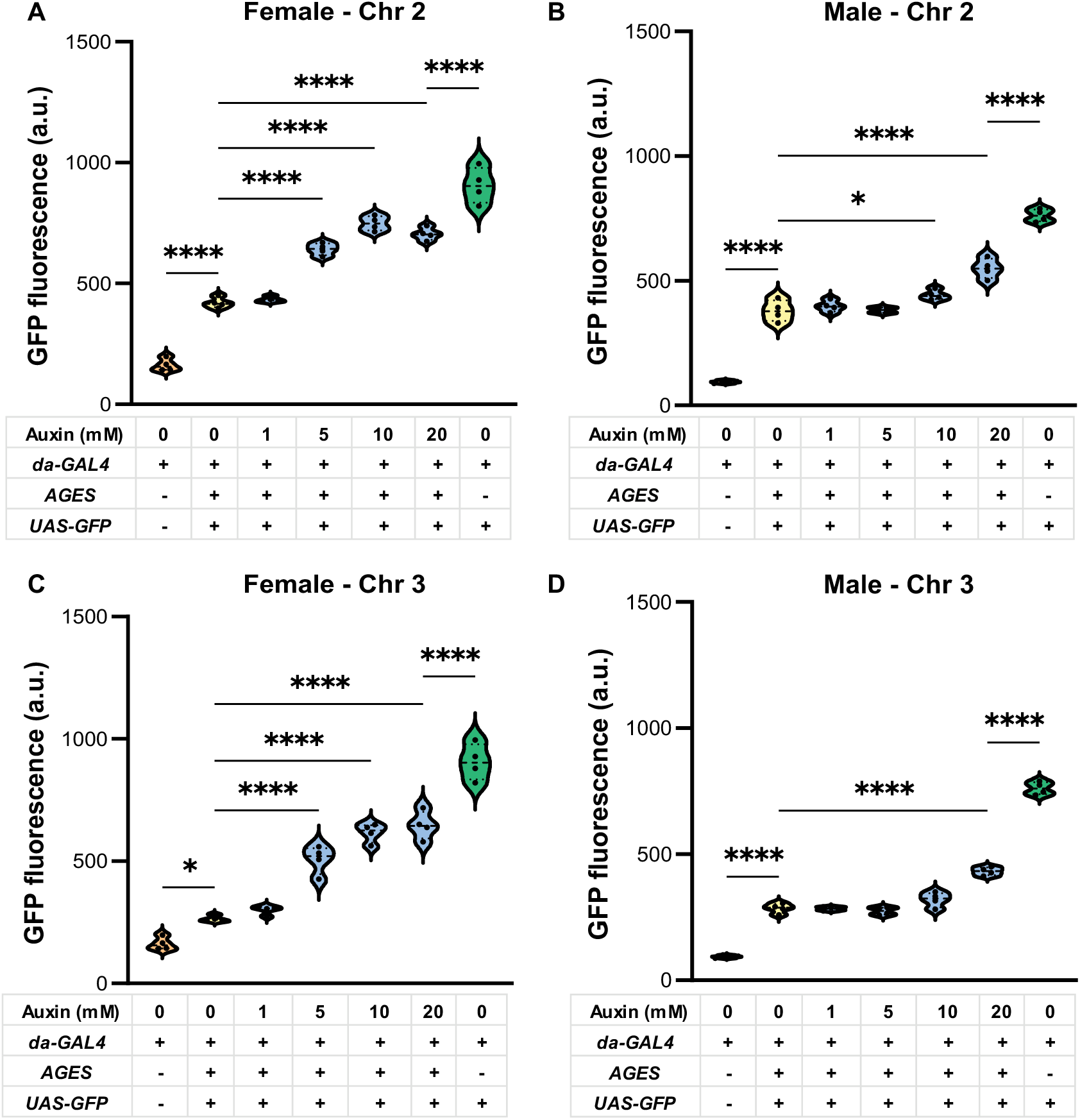
Transgene induction with AGES varies with sex and transgene insertion site. GFP quantification (expressed as arbitrary units, a.u.) was derived from fluorescence intensity readings of whole-body fly homogenates following a 7-day induction period on media containing different concentrations of auxin. Data are shown as female (A) and male (B) flies where *AGES* (*Tub84B-AtTIR1-T2A-AID-GAL80-AID*) was located on the second chromosome, or females (C) and males (D) where *AGES* was located on the third chromosome. Data are presented as violin plots (line at median) of n = 4 replicates per condition, each containing 3 flies per replicate. Note that the negative control (*da-GAL4* alone) and positive control (*da-GAL4>UAS-GFP* alone) groups are identical in A/C and B/D, since these samples were run in parallel. Statistical comparisons were performed using one-way ANOVA with subsequent Bonferroni-corrected multiple comparison tests where *, p < 0.05 and ****, p < 0.0001.

**Figure 2.**
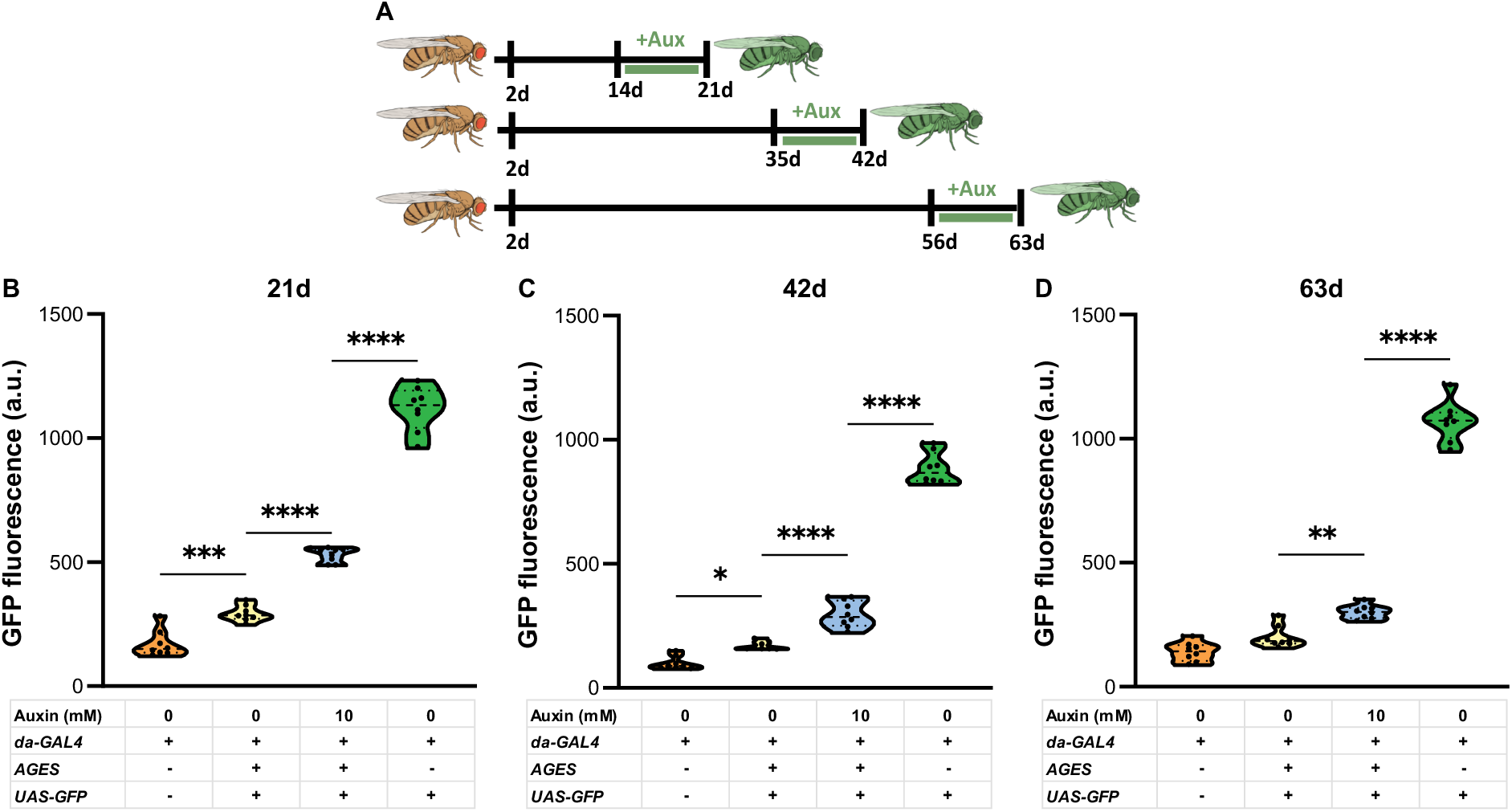
Transgene induction with the AGES system remains permissible at older ages. Schematic depicting the auxin feeding protocol which utilised differentially aged fly cohorts to assess GFP induction (A). GFP quantifications (expressed as arbitrary units, a.u.) were derived from fluorescence intensity readings of whole-body fly homogenates following a 7-day induction period on 10 mM auxin or control (0 mM) media. Data are shown for fly cohorts aged to 21d (B), 42d (C), and 63d (D) at the time of assay. Data are presented as violin plots (line at median) of n = 8 replicates per condition, each containing 3 flies per replicate. Statistical comparisons were performed using one-way ANOVA with subsequent Bonferroni-corrected multiple comparison tests where *, p < 0.05; **, p < 0.01; ***, p < 0.001; and ****, p < 0.0001.

**Figure 3.**
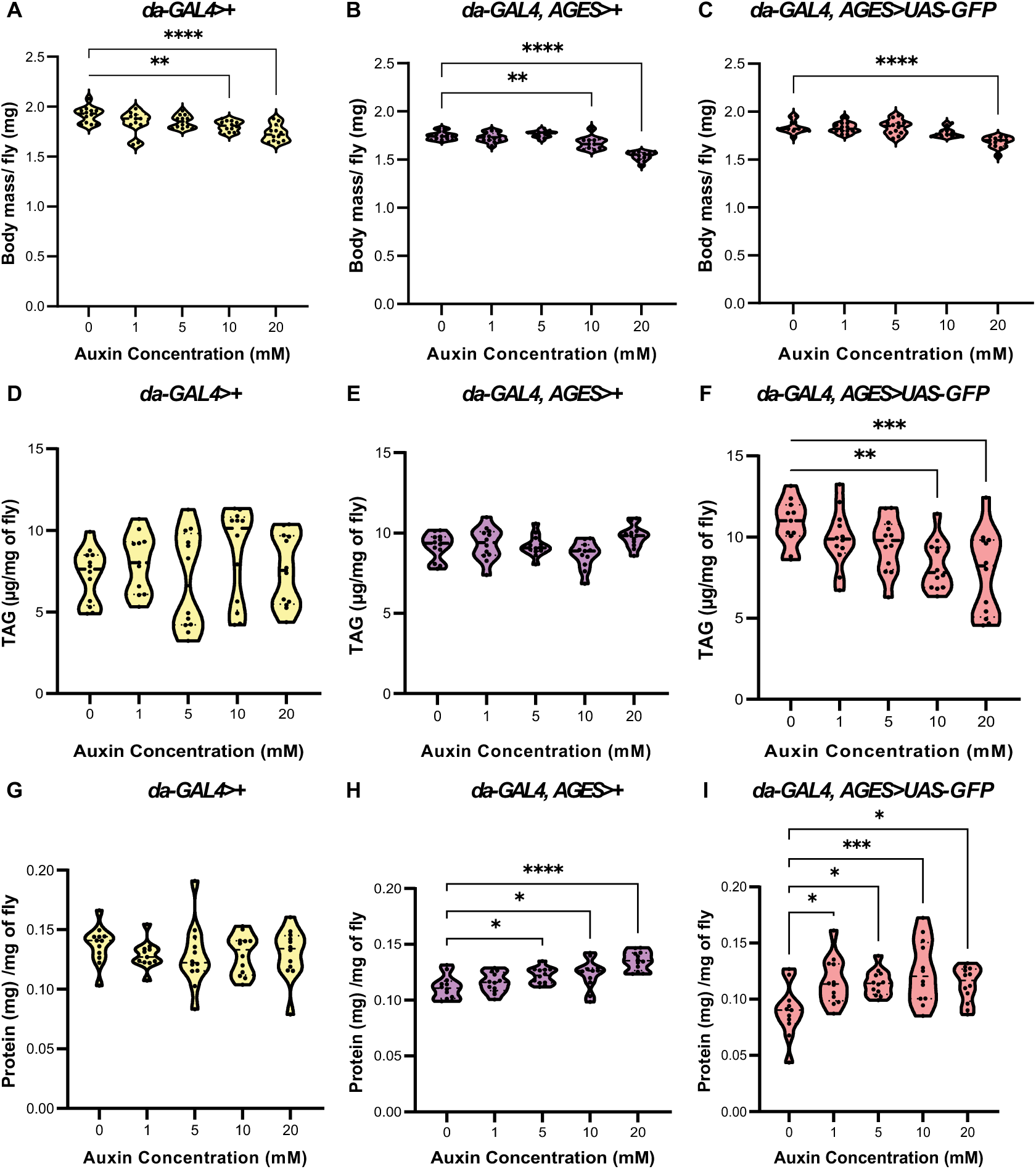
High-dose auxin reduces body mass independent of genotype, and alters TAG and protein levels in genotype-dependent ways. Whole body mass measurements are shown for *da-GAL4>+* (A), *da-GAL4,AGES>+* (B), and *da-GAL4,AGES>UAS-GFP* (C) female flies fed different concentrations of auxin for 7 days. Triacylglycerol (TAG) measurements are depicted for *da-GAL4>+* (D), *da-GAL4,AGES>+* (E), and *da-GAL4,AGES>UAS-GFP* (F) fly cohorts. Quantification of protein concentration are illustrated for *da-GAL4>+* (G), *da-GAL4,AGES>+* (H), and *da-GAL4,AGES>UAS-GFP* (I) groups. Data are described as per fly body weight (A-C) or normalised to per mg of fly for both TAG and protein values (D-F and G-I), where all metrics were derived from the same samples for a given genotype. Data are presented as violin plots (line at median) of n = 11-12 replicates per condition, each containing 5 flies per replicate. Statistical comparisons were performed using one-way ANOVA with subsequent Dunnett’s multiple comparison tests to corresponding 0 mM control conditions where *, p < 0.05; **, p < 0.01; ***, p < 0.001; and ****, p < 0.0001.

**Figure 4.**
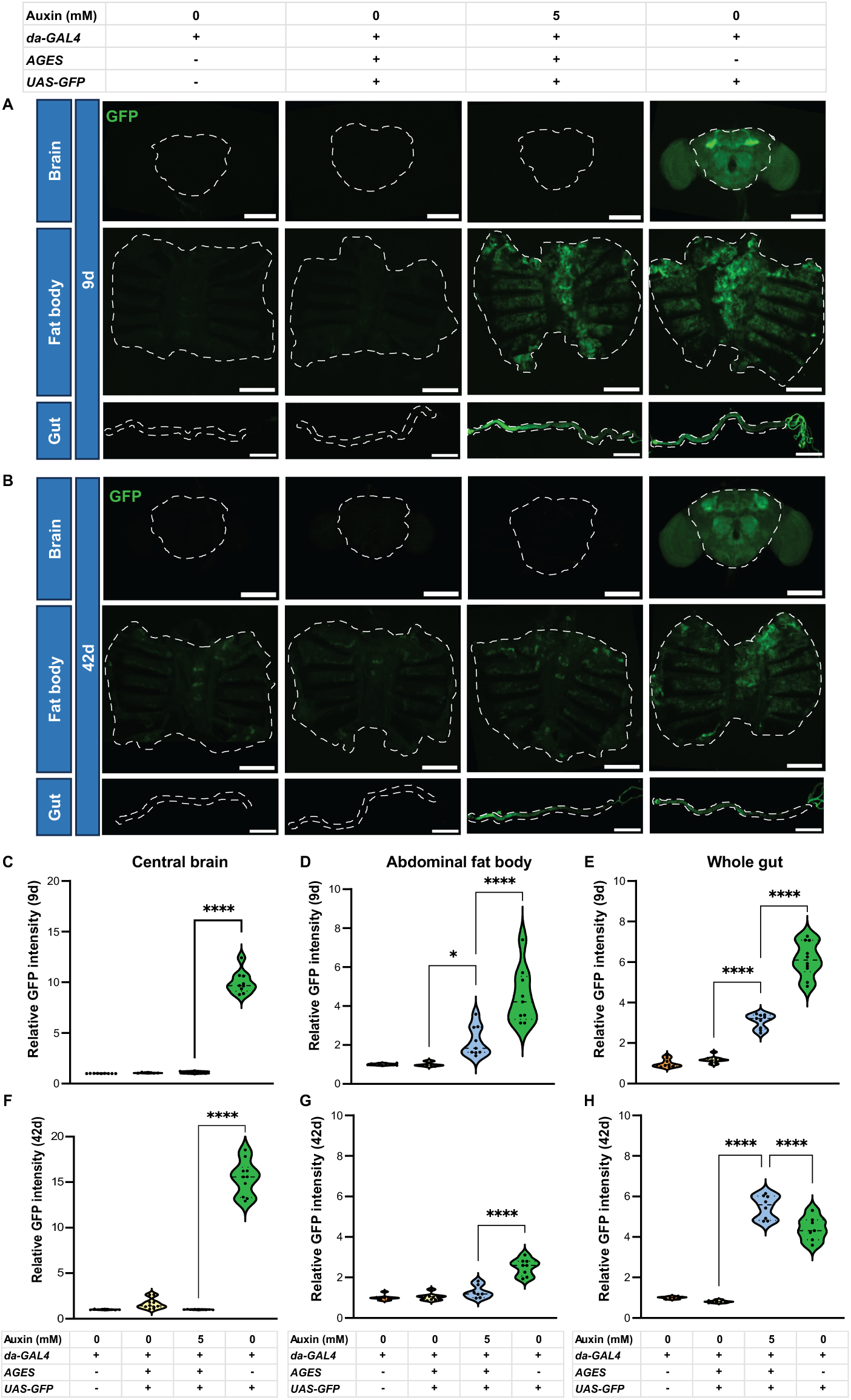
AGES shows tissue- and age-dependent expression patterns with the *da-GAL4* driver. Adult brain, abdominal fat body, and whole gut GFP channel images (green) are shown from dissected fly cohorts following a 7-day induction period on 5 mM auxin or control (0 mM) media, where flies were aged to 9d (A) and 42d (B) at the time of dissection. Dotted lines indicate traces performed on DAPI stains from the same images of the central brain, abdominal cuticle with attached adipose tissue and whole gut length. Corresponding mean GFP fluorescence intensity values (expressed relative to *da-GAL4*-alone-containing groups) are shown for the central brain (C), abdominal fat body (D) and whole gut (E) of 9d flies, alongside brain (F), fat body (G) and gut (H) measurements from 42d flies. Data are presented as violin plots (line at median) of n = 6-10 flies per condition. Statistical comparisons were performed using one-way ANOVA with subsequent Bonferroni-corrected multiple comparison tests where *, p < 0.05 and ****, p < 0.0001. Scale bars, 200 μm (brain), 500 μm (fat body) and 1 mm (gut).

### Whole-Fly GFP Quantification assays

For all GFP-based plate reader assays, flies were maintained on auxin or control food at varying concentrations and starting ages for 7 days as specified for each experiment. Flies (3 per replicate) were then homogenised in Eppendorf tubes containing 240 µL of 1X RIPA lysis buffer (diluted from a 10X stock (#20-188, Millipore) plus protease inhibitors (added from a 7X solution (#11836170001, Roche). Whole-fly homogenates were obtained by motorised homogenisation followed by centrifuging each sample at 4°C and 20,800 × *g* for 2 minutes.

80 µL of the resulting supernatant was added from each sample to a black, flat bottom 96 well plate (#237105, Fisher Scientific). Standard curves were generated by serially diluting the pooled homogenate of 10 positive control flies (ubiquitously expressing mCD8::GFP) in duplicate. Full plate adjustment was used to optimise the gain setting per run, and fluorescence intensity was measured (488 nm excitation; and 505 nm emission) in a Thermo MultiSkan FC plate reader. Fluorescence intensity values for each sample were then quantified relative to the standard curve values.

### Triacylglycerol (TAG) and BCA assays

For all TAG assays, flies were maintained on auxin or control food at varying concentrations from day 2 of adulthood for 7 days. Female flies were then divided into groups of 5 and weighed on a Thermo microbalance to an accuracy of .01 mg. Weighed females were then added to 2mL screw-cap tubes containing 1.4mm ceramic beads (#19-645-3, Cole-Parmer) with 125 µL of TET buffer (10mM Tris (#T6066, Sigma), 1mM EDTA (#10526383, Fisher Scientific), 0.1% (v/v) Triton-X-100 (#648466, Millipore)). Samples were homogenised via an Omni Bead Ruptor Elite bead mill homogeniser at 6.45 (m/s) for one minute and then immediately incubated at 72°C for 15 min in a heat block to inactivate lipases/other degradative enzymes. Sample tubes were then centrifuged at 4°C and 20,800 × *g* for 5 minutes before adding 2 µL of the supernatant from each sample or glycerol stock (1-0 µg/µL range) to a clear, flat bottom 96-well plate (Corning Life Sciences). 300 µL of TAG reagent (full constituents listed in Supplementary Table 1) was added to each well and absorbance measured at 540 nm in a Thermo MultiSkan FC plate reader. Triacylglycerol levels for each sample were then quantified relative to the standard curve values.

The BCA Protein Assay Kit (#23227, Pierce) was used to assay protein content from frozen sample homogenates as per the manufacturer’s instructions.

### Fly tissue dissections and imaging

Adult brains were dissected in PBT (1X phosphate-buffered saline (PBS) containing 0.5% Triton X-100) and fixed in 4% paraformaldehyde (PFA) in PBT for 20 minutes at room temperature. After fixation, samples were washed with PBT three times for 20 minutes at room temperature, incubated in Vectashield with DAPI (#H-1200-10, Vector Laboratories) overnight at 4 °C and subsequently mounted on slides. Whole guts were dissected in 1X PBS and fixed in 4% PFA in PBS for 30 minutes in a 6 well culture plate (Corning Life Sciences). Guts were then incubated in 1 mL PBT overnight at 4 °C. PBT was aspirated from sample wells and samples were washed an additional three times for 20 minutes each before being mounted in Vectashield with DAPI. For fat body dissections, gut, tubules and reproductive anatomy were removed from isolated fly abdomens, leaving abdominal cuticle with adipose tissues attached. Thereafter, samples were fixed, washed and mounted as for gut tissues above. All slides were stored at 4°C in the dark before imaging.

Images were acquired using a 5X objective on a Leica DM5500 B microscope via a K3M Leica camera and equipped with LASX (3.8.1.26810.1) software, where the optical settings were kept constant between samples of a given tissue. The Polygon selection tool in Fiji (ImageJ) (Schindelin *et al*., 2012) was used to trace regions of interest (ROI’s) which corresponded to either the central brain (chosen due to non-uniformity of the optic lobes from dissection), whole gut, or abdominal fat body, all of which were visualised/traced in the DAPI channel. GFP mean fluorescence intensity (MFI) per ROI was analysed using Fiji to obtain measurements for a total of 6-10 biological replicates for a given condition per tissue group.

### Quantitative real-time PCR (qPCR)

Total RNA was isolated from whole flies via TRIzol (#12034977, Invitrogen), as per the manufacturer’s instructions. RNA samples were then subjected to DNA digestion via ezDNase (#15585209, Invitrogen), followed by conversion to cDNA using oligo(dT) primers (#11685581, Invitrogen) and SuperScript IV Reverse Transcriptase (#15307696, Invitrogen). qPCR was performed using 2x QuantiNOVA SYBR Green PCR Master Mix (#208054, Qiagen) in a 7500 Fast Real-Time PCR machine. Relative quantities of transcripts were determined through the relative standard curve method, normalised to the geometric mean of *eIF1A* and *Tub84B* reference genes, which have demonstrated good stability in samples from *Drosophila* across ageing (Ling and Salvaterra, 2011). Primer sequences were as follows:

eIF1A_for: ATCAGCTCCGAGGATGACGC; eIF1A_rev:

GCCGAGACAGACGTTCCAGA; InR_total_for: CGGACGACCGTCAAAGGTTA;

InR_total_rev: GCCTTGACGAGAGGTTCAGT; InR_DN[K1405A]_for:

ATCGCGAGTGTGCCATTGC; InR_DN[K1405A]_rev:

GGAGCAAACACCGAGCAATC; InR_WT_[K1405]_for:

GATCGCGAGTGTGCCATTAA; InR_WT_[K1405]_rev:

GGAGCAAACACCGAGCAATC; Tub84B_for:

TGGGCCCGTCTGGACCACAA; Tub84B_rev:

TCGCCGTCACCGGAGTCCAT.

### Starvation and Oxidative Stress assays

For all stress assays, flies were maintained on auxin or control food at the described concentrations from day 2 of adulthood until 14 days of age, with an n of ∼150 flies for a given condition (density of 15 flies per vial). For starvation, flies were transferred to vials containing 1.5% (w/v) agarose (#16500-500, Invitrogen) to permit hydration but restrict nutritional value. Deaths and censors were scored two times per day, with flies transferred to fresh vials three times per week. For oxidative stress assays, flies were transferred to vials containing 5% (v/v) H_2_O_2_ (#216763, Merck) (constituted in 1.5% (w/v) agar and 5% (w/v) sucrose) and similarly transferred to fresh vials three times per week with deaths and censors scored two times per day.

### Lifespan Assays

Lifespan assays were set up as above, with an n of ∼150 flies for a given condition (density of 15 flies per vial). Flies were transferred to fresh vials three times per week where deaths and censors were recorded during each transfer. Microsoft Excel (scoring template available at http://piperlab.org/resources/) was used to calculate survival proportions after each transfer.

### Female fecundity assay

Fecundity assays were set up as above, with an n of ∼50 flies for a given condition (density of 5 flies per vial). Flies were transferred to fresh vials three times per week, and the number of eggs laid per vial over a 20-24h period was manually counted once weekly. Data are presented as the cumulative number of eggs laid per female per 24h over the entire measured period (1 through 5 weeks of age), calculated in Microsoft Excel (http://piperlab.org/resources/).

### Statistical Analysis

Statistical analyses were either carried out in Microsoft Excel or GraphPad Prism 10.6.1 with tests for a given experiment detailed in each figure legend. Exact lifespan and stress assay deaths and censors (corresponding to the final sample sizes) are detailed in full within the source data. For all statistical tests (specific tests listed in each figure legend), *p* < 0.05 was considered significant.

### Data Availability

Source data underlying all figures are available at https://doi.org/10.17632/rw4mvv8mk2.1

## Results

### Transgene induction via AGES demonstrates sex and insertion-site-specific variability

To investigate applicability of AGES in the same context as our longevity experiments, we characterised transgene induction in both sexes, from different transgene insertion sites, and across an auxin titration (∼0-20 mM) (as per McClure et al., 2022). Since we used the potassium salt of auxin, we added equimolar potassium chloride as a vehicle control throughout (see Methods). To assess transgene expression, we expressed membrane-bound GFP (*UAS-mCD8::GFP,* hereafter termed *UAS-GFP*) ubiquitously, via *da-GAL4* under control of the *Tub84B-AtTIR1-T2A-AID-GAL80-AID* transgene (hereafter termed *AGES*) inserted on either the second or third chromosome. We fed auxin from days 2-9 of adulthood, then quantified GFP fluorescence (**Figure 1**).

In both sexes and from both insertion sites, GFP fluorescence was significantly increased in the control-fed (0 mM auxin) *da-GAL4,AGES>UAS-GFP* flies (full genotypes *w^Dah^;AGES/+;da-GAL4,UAS-mCD8::GFP/+* or *w^Dah^;+;da-GAL4,UAS-mCD8::GFP/AGES*) compared to the negative control groups (containing the *da-GAL4* construct alone, full genotype *w^Dah^;+;da-GAL4/+*), suggesting a degree of GAL4 activity even in the absence of auxin feeding (**Figure 1A-D**). GFP fluorescence was increased by ≥5 mM auxin in females, regardless of AGES transgene insertion site (**Figure 1A and C**). In males bearing *AGES* on the second chromosome, we observed significant GFP induction only at 10 mM and 20 mM auxin, while males with *AGES* on the third chromosome exhibited increased GFP only at 20 mM auxin (**Figure 1B and D**). When *AGES* was present, GFP fluorescence did not reach the level of positive controls (*da-GAL4>UAS-GFP*, full genotype *w^Dah^;+;da-GAL4,UAS-mCD8::GFP/+*), even when using the highest auxin dose (**Figure 1A-D**). Together, these results suggest that transgene induction depend on sex and, in males, *AGES* insertion site; and that some residual GAL80 activity may occur in both sexes and from both transgene insertion sites.

In all subsequent experiments, we utilised female flies with the *AGES* construct on the third chromosome, owing to its strongly inducible expression at 5, 10, and 20 mM auxin as well as its lower leakiness at 0 mM (**Figure 1C**).

### AGES permits controlled transgene induction throughout the fly life course

We next asked whether the AGES system could induce transgene expression effectively and in a stable manner as flies age, given known age-associated changes in efficacy of alternative established systems (Barwell *et al*., 2017; Delandre, McMullen and Marshall, 2025). Again, using *da-GAL4,AGES>UAS-GFP* flies, we established young, middle-aged and old cohorts (14d, 35d, and 56d, respectively), then fed auxin (10 mM) for 7 days, and assessed GFP induction (**Figure 2A**). At all ages, we found that flies fed auxin had significantly increased GFP expression compared to 0 mM control conditions (**Figure 2B-D**). At 21d, auxin treatment produced a 1.8-fold increase in mean GFP signal compared to the 0 mM group (**Figure 2B**), whereas at 42d (**Figure 2C**) and 63d (**Figure 2D**) this fold increase was 1.7 and 1.5 respectively. However, these induction levels were still significantly less than in positive control (*da-GAL4>UAS-GFP)* flies (at 21d, the auxin-fed mean GFP signal was only 47%; at 35d only 33%; and at 63d, only 28%). At both 21d and 42d (but not 63d), fluorescence in flies bearing *UAS-GFP* was greater than negative controls (*da-GAL4*/+) even in the absence of auxin (**Figure 2B-D**), indicating that AGES does not completely repress GAL4 activity at these two younger timepoints. Altogether these results suggest that AGES can be used to conditionally induce transgene expression across ages, although induction does not fully recapitulate expression without *AGES* transgenes, and residual transgene expression is possible in youth and middle age even in the absence of induction by auxin.

### Auxin exerts genotype-dependent and dose-dependent effects on metabolic indices

Auxin can alter fly metabolism, for example causing a decrease in triacylglycerol (TAG) levels (Fleck *et al*., 2024). We therefore tested whether we recapitulated metabolic effects in flies treated with auxin (0-20 mM) for 7 days (days 2-9 of adulthood). We measured body mass, TAG, and protein levels in flies expressing *da-GAL4* alone (full genotype *w^Dah^;+;da-GAL4/+*), *da-GAL4*,*AGES* alone (full genotype *w^Dah^;+;da-GAL4/AGES*) and *da-GAL4,AGES>UAS-GFP* (full genotype *w^Dah^;+;da-GAL4,UAS-mCD8::GFP/AGES*). These conditions allowed us to test the effects of auxin in flies lacking *AGES* transgenes, as well as the effect of auxin-induced GAL4 activity in flies with or without a *UAS-trangene*.

In all genotypes, body mass was reduced by ≥10mM auxin (**Figure 3A-C**). TAG levels were not affected by auxin in either *da-GAL4* or *da-GAL4*,*AGES* (**Figure 3D and E**), but ≥10 mM auxin decreased TAG in *da-GAL4,AGES>UAS-GFP* flies (**Figure 3F**). Protein levels were not affected by auxin in *da-GAL4* (**Figure 3G**), but were increased by ≥5 mM auxin in *da-GAL4*,*AGES* and ≥1 mM auxin in *da-GAL4,AGES>UAS-GFP* (**Figure 3H and I**). Taken together, these results indicate that high dose auxin affects body mass regardless of the presence of AGES, and that there are further AGES-dependent and/or transgene/GFP-dependent effects on TAG and protein levels. In subsequent experiments, we therefore used 5 mM auxin, aiming to minimise off-target metabolic effects of auxin while still achieving sufficient transgene induction (**Figure 1C**).

### Ubiquitous transgene induction via AGES displays tissue-specific expression variability

Given the reliance on homogenised whole-fly extracts within our fluorescence intensity assays which mask tissue-specific expression levels, we next investigated whether, AGES can induce transgene expression effectively across multiple organs in both young and older flies. To verify this, we fed *da-GAL4, AGES>UAS-GFP* flies auxin (5 mM) for 7 days starting at day 2 and day 35 of adulthood, representing a young (9d) and older (42d) timepoint respectively, when we examined GFP expression in the brain, whole gut, and abdominal fat body tissues following dissection (**Figure 4**). Notably, these tissues cover a range of expression strengths for the *da-GAL4* driver, which, while strongly expressed in many peripheral tissues, has previously been reported to drive expression at lower levels in the brain than some other ubiquitous or neuron-specific drivers (Legan *et al*., 2008).

Within our experiments, auxin treatment (5 mM) did not increase GFP fluorescence in the adult brain in either young or old flies (**Figure 4A-C and F**). However, auxin treatment increased GFP fluorescence in the fat body of young but not old flies (**Figure 4A and D**) (**Figure 4B and G**). In youth, this fat body induction was 49% of that displayed by the positive control (*da-GAL4*>*UAS-GFP*) condition (**Figure 4D**). In the gut, auxin increased GFP signal in old and young flies (**Figure 4A, B, E and H**). At 9d, induction in guts was equivalent to 50% of the positive control (**Figure 4E**), but at 42d, unexpectedly 24% higher in the auxin-induced group than in the positive control, **Figure 4H**). Finally, across all tissues and both timepoints, we observed negligible transgene leakiness by these assays (**Figure 4A-H**). These results suggest the presence of tissue- and age-specific differences in expression with the AGES system, but they also demonstrate that in some tissues such as the gut, AGES permits strong transgene expression even at older ages.

### AGES-directed expression of InR^DN^ only partially reproduces known effects of reduced IIS on lifespan and fecundity

We next tested whether AGES recapitulated the extension of healthy lifespan associated with reduced insulin/insulin-like signalling (IIS), an evolutionarily-conserved phenotype (Mathew, Pal Bhadra and Bhadra, 2017). Using AGES, we expressed a dominant-negative insulin receptor variant (*UAS-InR^DN^*, encoding the kinase-dead InR^K1405A^ variant) throughout adulthood, mimicking an established lifespan-extending manipulation using GeneSwitch (Slack et al., 2011). We confirmed by qPCR that auxin treatment (5 mM from day 2 to 9 of adulthood) induced *InR^DN^* expression in *da-GAL4,AGES>UAS-InR^DN^*flies (full genotype *w^Dah^;UAS-InR^DN^/+;da-GAL4,AGES/+*) using primers that amplify either wild-type-specific, DN-specific, or both sequences (**Figure 5A**). We found that auxin treatment substantially induced expression of *InR^DN^*, with a small but significant elevation in wild-type *InR* mRNA level (**Figure 5A**), potentially representing compensatory up-regulation of *InR* as reported previously for GeneSwitch-driven *InR^DN^* expression in flies (Graze *et al*., 2018).

**Figure 5.**
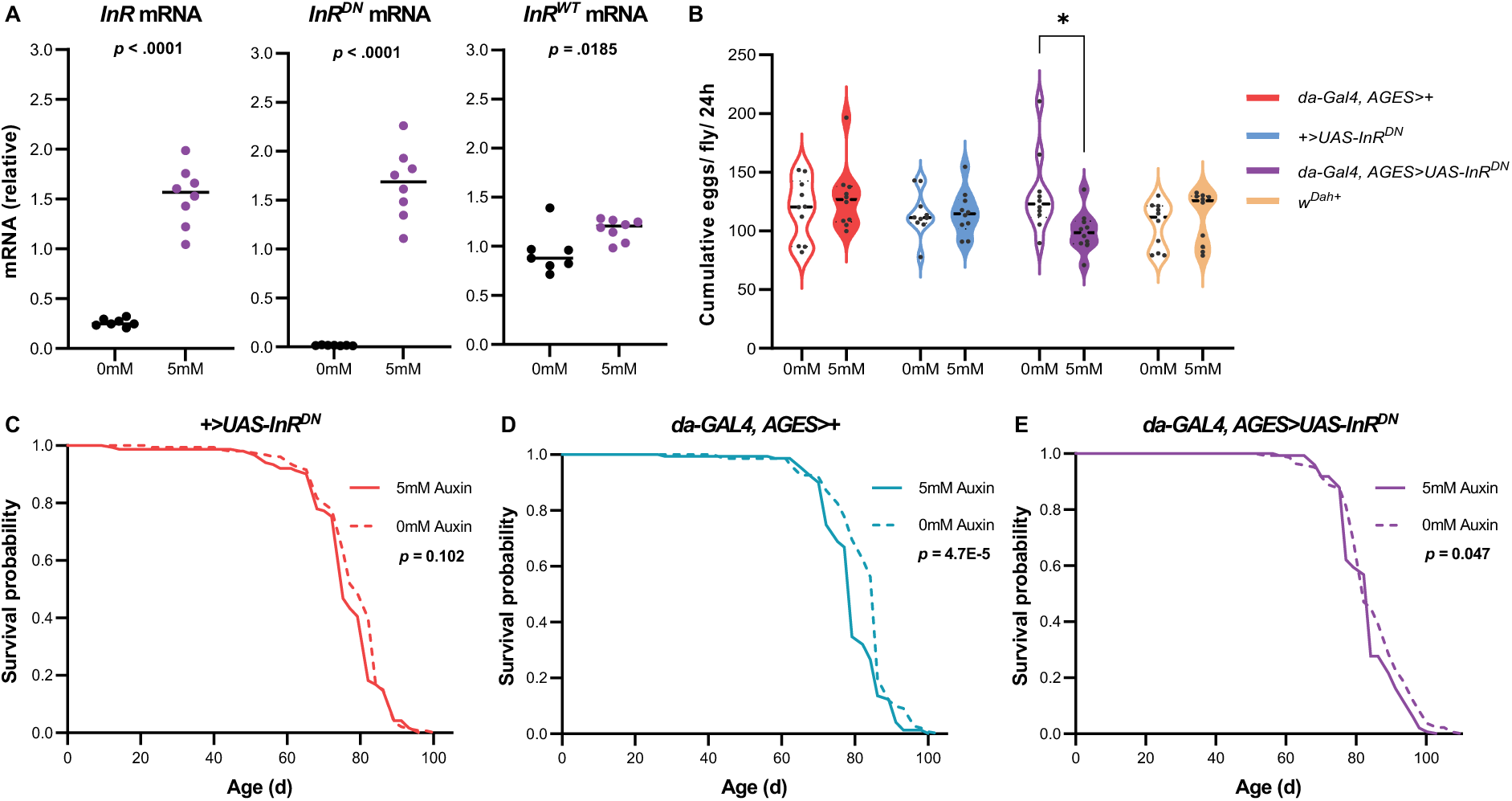
AGES-driven expression of *InR^DN^* reduces fecundity but has unclear effects on longevity. (A) Whole-fly mRNA levels for total *InR* (primers outside of K1405 sequence*), InR^DN^* (primers for K1405A) and *InR^WT^* (primers for wild-type K1405) are shown for *da-GAL4,AGES>UAS-InR^DN^* female flies following a 7-day induction period on 5 mM auxin or control (0 mM) media from day 2 to day 9 of adulthood prior to collection for qPCR. Data are presented as scatter plots (line at median) of n = 7 (0mM) or 8 (5mM) replicates per condition, each containing 2 flies per replicate. (B) Quantification of egg-laying over a 34-day period is illustrated for female flies of the indicated genotypes, treated with 5 mM auxin or control (0 mM) media. Data are depicted as the cumulative number of eggs laid per female per 24h, and presented as violin plots (line at median) of n = 10 replicates per condition, each containing 5 flies per replicate. (C-E) Lifespan curves are depicted for *+>UAS-InR^DN^* (C), *da-GAL4,AGES>+* (D), and *da-GAL4, AGES>UAS-InR^DN^* (E) female flies following continual 5 mM auxin or 0 mM treatment from day 2 of adulthood. For all survival data, n > 140 deaths were counted per condition, with survival curves and p-values derived via Log-Rank test analyses. For qPCR and fecundity data, statistical comparisons were performed using unpaired t-tests and a two-way ANOVA with Bonferroni’s multiple comparison test respectively, where *, p < 0.05.

Given that increased longevity mediated through lower IIS is frequently, but not always, accompanied by a reduction in fecundity (Flatt, 2011; Ibrahim *et al*., 2026), we quantified fecundity impact of *InR^DN^*expression under control of *AGES* (via weekly egg counts over a 34-day period). Consistent with previous work using GeneSwitch (Slack *et al*., 2011), we found that *UAS-InR^DN^* induction by auxin feeding reduced egg laying, whereas fecundity of control genotypes was not affected by auxin (**Figure 5B**).

In parallel to fecundity assays, we also measured lifespan of flies fed auxin continuously from day 2 of adulthood. Here, we observed complex, genotype-dependent effects of auxin. First, we found no significant difference in the lifespan of auxin-treated *+>UAS-InR^DN^* flies (full genotype *w^Dah^;UAS-InR^DN^/+;+*) (**Figure 5C**). However, auxin shortened lifespan of *da-GAL4,AGES>+* flies (full genotype *w^Dah^;+;da-GAL4,AGES/+*) (**Figure 5D**: median -8.3%). In *da-GAL4,AGES>UAS-InR^DN^*flies, auxin led to a significant but complex and relatively modest change in longevity (**Figure 5E**: median lifespan +3.2%, but with reduced survival at other ages). Taken together, these results suggest that auxin does not affect fly lifespan in the absence of the *AGES* construct, but de-repressing GAL4 by feeding auxin in the presence of *AGES* can shorten lifespan. When transgenes are recombined to enable auxin-dependent *InR^DN^* expression, AGES can recapitulate established effects of *InR^DN^* expression (decreased fecundity and compensatory up-regulation of wild-type *InR* for instance), but the interpretation of lifespan data is challenging here given the effects of auxin in control genotypes.

### Auxin impairs starvation and oxidative stress resistance

*UAS-InR^DN^* induction in adults via the GeneSwitch system increases resistance to oxidative stress and starvation stress (Slack *et al*., 2011; Bolukbasi *et al*., 2017). However, responses to these stresses are related to lipid metabolism, which has been shown to be affected by auxin/AGES (Fleck *et al*., 2024). We therefore tested whether *InR^DN^* induction by AGES recapitulated the stress resistance phenotypes shown previously following *InR^DN^* induction by GeneSwitch. We fed flies auxin or control media from days 2-14 of adulthood, then transferred them to 5% (v/v) H_2_O_2_ - containing food for oxidative stress (or agarose-only media for starvation) and measured survival in +>*UAS-InR^DN^*alone, *da-GAL4*,*AGES*>+, and *da-GAL4*,AGES>*UAS-InR^DN^*flies. We found that auxin universally reduced both oxidative stress resistance and starvation resistance across all genotypes (**Figure 6A-F**), suggesting a detrimental effect of auxin treatment independent of the presence or absence of *AGES*- or *InR^DN^*-encoding transgenes.

**Figure 6.**
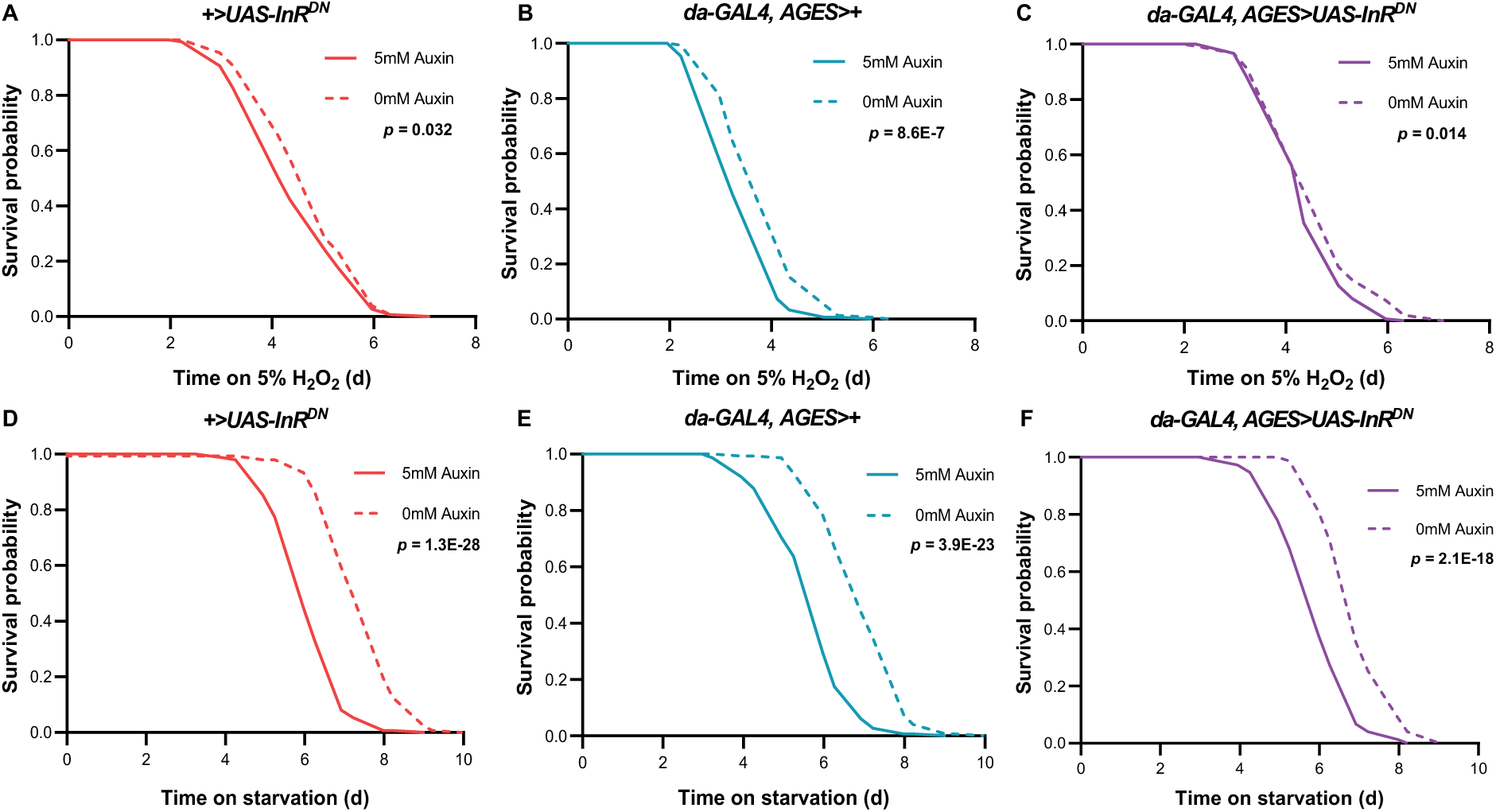
Auxin treatment reduces oxidative stress resistance and starvation resistance. (A-C) Survival curves are depicted for *+>UAS-InR^DN^* (A), *da-GAL4,AGES>+* (B), and *da-GAL4,AGES>UAS-InR^DN^* (C) female flies under H_2_O_2_ -based oxidative stress following 5 mM auxin or control treatment (0 mM) for 12 days. (D-F) Survival curves are shown for *+>UAS-InR^DN^* (D), *da-GAL4,AGES>+* (E), and *da-GAL4,AGES>UAS-InR^DN^* (F) female flies under starvation following the same 5 mM auxin treatment regimen. For all survival data, n > 145 deaths were counted per condition, with survival curves and p-values derived via Log-Rank test analyses.

## Discussion

Since its establishment, the spatiotemporal gene expression system AGES has been used to investigate questions in fields from immunity to neuroscience in *Drosophila* (Richhariya *et al*., 2023; Corke *et al*., 2025; Sobrido-Cameán *et al*., 2025; Kuklinski *et al*., 2026). Our study tests the additional application of ageing biology. Here, we have characterised the capability of the system to permit transgene induction across conditions typical for *Drosophila* ageing and longevity experiments, where we observed that transgene expression depends on auxin dose, age, sex, and tissue. Conversely, we have further confirmed that auxin and/or activation of the AGES machinery exert complex effects on fly metabolism which need to be considered in designing studies assessing longevity or other indices of organismal health.

When characterising feeding conditions commonly used for ageing experiments, we found that transgene induction was most optimal in female flies, fed in the 5-20 mM dose range of auxin, with the *AGES* transgenic construct inserted on the third chromosome (**Figure 1C**). Regarding sex-dependent differences, it is likely that these are, at least in part, underpinned by known differences in food consumption, where females ingest greater amounts driven by metabolic demands for oogenesis (Wong *et al*., 2009; Vargas *et al*., 2010). Notably, however, even at the highest auxin dose, transgene induction never reached the levels displayed by positive control flies lacking the *AGES* construct (*da-GAL4>UAS-GFP*), an observation in agreement with the original study describing the AGES system (McClure *et al*., 2022). Here, it is worth noting that the positive control flies will have been expressing GFP throughout development and adulthood, so the comparison to adult-onset expression via AGES would not necessarily be expected to achieve fully equal expression levels. Other work has also shown that parallel gene expression systems like TARGET can achieve higher transgene expression than that achievable with AGES, despite AGES demonstrating significantly less apparent transgene leakiness (Hawley, Roberts and Fitzsimons, 2023).

We also demonstrated that throughout the fly life course transgene induction remains feasible with AGES, albeit with decreased magnitudes with increasing age (**Figure 2**). Therefore, AGES may remain a viable option to permit the study of transgene-based manipulations in the aged fly following acute induction periods. However, given our testing of a single ubiquitous driver (*da-GAL4*), along with recognised GAL4-driver-specific differences in activity with age in other systems like TARGET (Delandre, McMullen and Marshall, 2025), driver selection should be carefully vetted when utilising AGES for ageing experiments in future studies. Our results agree with prior work that a 5 mM auxin dose is sufficient to elicit transgene induction in fat body tissue (McClure *et al*., 2022), and we further demonstrate this in the gut (**Figure 4A and E**). However, unlike prior observations at 10 mM auxin (McClure *et al*., 2022), we found that 5 mM auxin could not induce transgene expression in the brain of flies, regardless of age (**Figure 4A-C and F**). This could reflect either limitations of our chosen concentration of auxin (5 mM) in crossing the blood-brain barrier, or the relatively low expression of *da-GAL4* within the nervous system (Legan *et al*., 2008). Within the fat body, our results suggest that only younger (but not older) flies can achieve significant transgene expression (**Figure 4A-B, D, and G**), potentially due to age-related decreases in feeding rates (Carey et al., 2006). Within the gut, we observed relatively strong transgene induction at young ages and a surprising level of induction in aged flies that exceeded that of the positive control (**Figure 4B and H**). While the mechanism underlying the higher than positive control expression is unclear, it is possible that positive control flies develop long-term adaptive responses to GAL4 and/or GFP expression, for example endogenous turnover mechanisms that could ameliorate established negative consequences of constitutive GFP expression on physiology (Mawhinney and Staveley, 2011). With increasing age, female flies also exhibit a loss of gut barrier integrity (Rera, Clark and Walker, 2012; Regan *et al*., 2016) that may also underlie the age-specific effects seen here.

Our study also corroborates previous evidence suggesting that auxin feeding exerts detrimental effects on flies independent of genotype. For example, we observed metabolic impacts of auxin doses 10 mM and greater in our feeding regimen, consistent with previous evidence (Fleck *et al*., 2024). In our hands, high dose auxin supplementation (10-20 mM range) demonstrated significant body mass decreases (3-12% difference in mean) (**Figure 3A-C**) independent of fly genotype, potentially driven by decreased feeding events in auxin-containing media at this concentration range, as previously observed (Fleck *et al*., 2024). Even with no difference in the total mass of flies upon treatment at a lower dose of 5 mM auxin (**Figure 3A-C**), we still observed a stark reduction in stress-related survival for all genotypes of auxin-fed flies under starvation or oxidative stress (**Figure 6**), which suggests that auxin exerts detrimental effects even in flies where body mass remains constant. These effects may be driven by stress responses mounted against auxin itself, as prior evidence has demonstrated that 10mM auxin feeding induces transcriptomic signatures akin to that of a xenobiotic response with upregulation of detoxification-related transcripts (Fleck *et al*., 2024). It remains possible that a similar physiological reprogramming is apparent at 5 mM too, which in turn impacts stress survival capabilities independently of overall mass differences.

In addition, our study has identified detrimental effects of activating the AGES machinery independent of auxin feeding *per se*. For example, we observed subtle changes in protein levels (but not TAG levels) in auxin-fed flies that possess the *AGES* transgenes, but we did not observe similar effects in flies lacking the *AGES* transgenes (**Figure 3G-H**). Here, our findings differ somewhat from those of (Fleck *et al*., 2024), who observed a >50% decrease in TAG/protein ratios for flies of multiple genotypes exposed to auxin. These differences in the nature and magnitude of metabolic effects may be due to distinctions in auxin feeding regimens (e.g. pre-starved and fed on liquid auxin-containing food in previous studies, versus constitution of auxin into fly media here), the period of auxin exposure, and/or potential fly strain-specific effects. As an additional example where activating the AGES machinery was detrimental in our hands, we observed shortening of lifespan with auxin feeding in *da-GAL4, AGES* flies but not in *UAS-InR^DN^* flies (**Figure 5C-D**). These findings could be explained by a detrimental effect of GAL4 activity itself, consistent with previous reports (Dravecz *et al*., 2022); or, alternatively, activation of the TIR1 enzyme itself may exert additional effects that shorten lifespan. In either case, our findings support the need for appropriate controls (e.g. driver-alone and responder-alone) in future work using the AGES system.

As a proof of principle for AGES as a viable alternative system for ageing research, our study also attempted to recapitulate a well-known pro-longevity phenotype frequently observed upon reduction of IIS via expression of InR^DN^ (Ikeya *et al*., 2009; Slack *et al*., 2011; Bolukbasi *et al*., 2017). Unfortunately, our recapitulation here remained limited, with reduced resistance to environmental stressors in flies treated with auxin regardless of genotype (**Figure 6**), and a shortening of overall lifespan when *da-GAL4, AGES* flies were fed auxin (**Figure 5**). These results make it difficult to interpret whether InR^DN^ expression affected lifespan in this context, and serve as a cautionary tale for using the AGES system in lifespan studies, at least when using the *da-GAL4* driver. In summary, our results suggest that researchers using flies to study ageing should broadly consider the potential trade-offs between metabolic effects associated with auxin, the bioavailability of auxin at tissues of interest with their selected dose, and potential differences in induction attainable across sexes and *AGES* transgene insertion sites, when deciding whether AGES may be best suited for their given investigation.

Despite the limitations of the AGES system we describe above, our findings also leave room for the system to be a useful tool in certain domains of ageing research. For example, while our studies here have used a single GAL4 driver that expresses throughout fly tissues, it would be informative to evaluate whether AGES can recapitulate other lifespan and healthspan phenotypes upon reduced IIS with tissue-specific drivers that may not exert the same magnitude of systemic effects. In addition, our ubiquitous driver system was still able to recapitulate the reduced fecundity phenotype frequently described with lowered IIS (**Figure 5B**) without any change in fecundity in control genotypes, consistent with previous findings describing auxin’s lack of impact on oogenesis (Fleck *et al*., 2024). Taken together, these results suggest that AGES may be well suited for phenotypes related to egg development and fecundity, despite its impact on some other phenotypes.

Finally, we should also acknowledge several limitations within our study. Here we prioritised the use of female flies due to our observation that males show greater leakiness in transgene expression and lower levels of transgene induction upon auxin feeding (**Figure 1**). Our studies therefore miss potential sex-dependent differences in lifespan assays and metabolic indices, in contrast to previous work which demonstrated sex-specific feeding behaviours in contexts of auxin exposure (Fleck *et al*., 2024). As an additional limitation, whilst we used KCl supplementation in control conditions to control for the effects of potassium derived from the K-NAA (auxin) compound, we did not fully control for potential confounding factors introduced by dietary chloride on fly physiology. We have also focused on a single ubiquitous driver (*da-GAL4*) and a single pro-longevity intervention (InR^DN^ expression) that, while considered a classical pro-longevity intervention, also shows genotype-dependent effects on its modulation of longevity (Ibrahim *et al*., 2026). Lastly, whilst we utilised a genetic background strain (*w^Dah^*), diet, and chemical-inducing feeding regimen that are frequently used in ageing studies (Ziehm, Piper and Thornton, 2013; Piper and Partridge, 2016), we cannot rule out strain-specific and diet-dependent differences from our results that may differ from other experimental procedures that utilise the AGES system.

Overall, our findings demonstrate that AGES remains a pertinent tool to modulate gene expression, even in the aged fly. However, our results also serve as a cautionary tale for the application of AGES when studying aspects of healthspan and longevity, primarily due to confounding effects of auxin and/or activation of the AGES machinery on metabolism. This caution aligns with previously described alterations in feeding behaviour upon auxin supplementation (Fleck *et al*., 2024). Here, we suggest that future studies using the AGES system would benefit from the inclusion of rigorous control genotypes (e.g. auxin-treated driver-alone and responder-alone controls). As an attractive future approach that may avoid some of these confounding effects, proposed alternate variants of the TIR1 receptor have the potential for efficacy at significantly lower concentrations of alternative auxins (McClure *et al*., 2022; Zhang *et al*., 2022). These systems may therefore exert reduced off-target effects on metabolism, thus providing additional opportunities for the use of AGES in future ageing studies within *Drosophila*.

## Acknowledgements

We thank Tracy Lamont, Chloe Buchanan, and the rest of the Glasgow Academic Support Unit (ASU) team for their tireless and professional work to prepare the fly media needed to carry out these experiments to a high standard. We further thank Jonathan Booth for experimental support/advice and the Sanz Montero lab for equipment sharing throughout this project. We also gratefully acknowledge Colin McClure (Queens University Belfast) for sharing *Drosophila* AGES stocks and advice on the AGES system, and Tony Southall (Imperial College London) for insightful comments and suggestions on this manuscript. This work was supported by the Wellcome Trust [301529/Z/23/Z]. AD was supported by a UKRI Future Leaders Fellowship [MR/Y019660/1].

## Author Contributions

T.M. and N.W. conceptualised the study; T.M., K.G., A.J.D. and N.W. designed the methodology; T.M., K.G., A.J.D. and N.W. conducted the investigation; T.M. and N.W. conducted formal analysis; T.M. wrote the original draft of the manuscript; T.M., K.G., A.J.D. and N.W. wrote, reviewed, and edited the manuscript; T.M. and N.W. visualised the study; N.W. supervised the study; N.W. acquired funding.

## Open Access Statement

For the purpose of open access, the authors have applied a Creative Commons Attribution (CC BY) licence to any Author Accepted Manuscript version arising from this submission.

## Supplementary Material

**Supplementary Table 1.** Full constituents of Triacylglycerol (TAG) reagent. Quantities and corresponding product numbers are listed for each component required to make up the final TAG reagent used throughout experiments.

| Component | Quantity | Product number | Manufacturer |
| --- | --- | --- | --- |
| 50 mM potassium phosphate buffer pH 7.2 | 5 mL | P0662 & 1048711000 | Sigma |
| Lipase (150 U/mL) | 0.15 mL | 437707 | Sigma |
| Glycerol kinase (0.05 U/mL) | 0.02 mL | G6142 | Sigma |
| L-α-glycerol-phosphate-oxidase (5 U/mL) | 1 mL | G9888 | Sigma |
| Horseradish peroxidase (0.5 U/mL) | 0.1 mL | P8250 | Sigma |
| 0.7 mM ATP | 1 mL | A2383 | Sigma |
| 0.3 mM 4-aminophenozone | 1 mL | A4382 | Sigma |
| 3 μM potassium ferricyanide | 1 mL | 455989 | Sigma |
| 0.6 mM MgCl<sub>2</sub> | 0.1 mL | M8266 | Sigma |
| 1.7 mM 3,5-dichloro-2-hydroxybenzenesulfonic acid (DHSA) | 1 mL | D4645 | Sigma |
| Triton X-100 | 0.01 mL | 648466 | Millipore |
| H<sub>2</sub>O | 89.62 mL | N/A | N/A |

## References

Barwell, T. et al. (2017) “Regulating the UAS/GAL4 system in adult Drosophila with Tet-off GAL80 transgenes,” PeerJ, 5, p. e4167. Available at: 10.7717/peerj.4167.

Bass, T.M. et al. (2007) “Optimization of Dietary Restriction Protocols in Drosophila,” The Journals of Gerontology: Series A, 62(10), pp. 1071–1081. Available at: 10.1093/gerona/62.10.1071.

Biteau, B., Hochmuth, C.E. and Jasper, H. (2008) “JNK Activity in Somatic Stem Cells Causes Loss of Tissue Homeostasis in the Aging Drosophila Gut,” Cell Stem Cell, 3(4), pp. 442–455. Available at: 10.1016/j.stem.2008.07.024.

Bolukbasi, E. et al. (2017) “Intestinal Fork Head Regulates Nutrient Absorption and Promotes Longevity,” Cell Reports, 21(3), pp. 641–653. Available at: 10.1016/j.celrep.2017.09.042.

Bolukbasi, E. et al. (2021) “Cell type-specific modulation of healthspan by Forkhead family transcription factors in the nervous system,” Proceedings of the National Academy of Sciences, 118(8). Available at: 10.1073/pnas.2011491118.

Brand, A.H. and Perrimon, N. (1993) “Targeted gene expression as a means of altering cell fates and generating dominant phenotypes,” Development, 118(2), pp. 401–415. Available at: 10.1242/dev.118.2.401.

Carey, J. et al. (2006) “Age-specific and lifetime behavior patterns in Drosophila melanogaster and the Mediterranean fruit fly, Ceratitis capitata,” Experimental Gerontology, 41(1), pp. 93–97. Available at: 10.1016/j.exger.2005.09.014.

Clancy, D.J. et al. (2001) “Extension of Life-Span by Loss of CHICO, a Drosophila Insulin Receptor Substrate Protein,” Science, 292(5514), pp. 104–106. Available at: 10.1126/science.1057991.

Corke, J. et al. (2025) “Maturation of GABAergic signalling times the opening of a critical period in Drosophila melanogaster,” Scientific Reports, 15(1), p. 40285. Available at: 10.1038/s41598-025-24116-2.

Delandre, C., McMullen, J.P.D. and Marshall, O.J. (2025) “Dynamic changes in neuronal and glial GAL4 driver expression during Drosophila aging,” GENETICS, 229(3). Available at: 10.1093/genetics/iyaf014.

Dravecz, N. et al. (2022) “Reduced Insulin Signaling Targeted to Serotonergic Neurons but Not Other Neuronal Subtypes Extends Lifespan in Drosophila melanogaster,” Frontiers in Aging Neuroscience, 14. Available at: 10.3389/fnagi.2022.893444.

Flatt, T. (2011) “Survival costs of reproduction in Drosophila,” Experimental Gerontology, 46(5), pp. 369–375. Available at: 10.1016/j.exger.2010.10.008.

Fleck, S.A. et al. (2024) “Auxin exposure disrupts feeding behavior and fatty acid metabolism in adult Drosophila,” eLife, 12. Available at: 10.7554/eLife.91953.

Graze, R.M. et al. (2018) “Perturbation of IIS/TOR signaling alters the landscape of sex-differential gene expression in Drosophila,” BMC Genomics, 19(1), p. 893. Available at: 10.1186/s12864-018-5308-3.

Hawley, H.R., Roberts, C.J. and Fitzsimons, H.L. (2023) “Comparison of neuronal GAL4 drivers along with the AGES (auxin-inducible gene expression system) and TARGET (temporal and regional gene expression targeting) systems for fine tuning of neuronal gene expression in Drosophila,” Micropublication Biology, pp. 10–17912.

He, Y. and Jasper, H. (2014) “Studying aging in Drosophila,” Methods, 68(1), pp. 129–133. Available at: 10.1016/j.ymeth.2014.04.008.

Ibrahim, R. et al. (2026) “Lifespan and Fecundity Impacts of Reduced Insulin Signalling Can Be Directed by Mito-Nuclear Epistasis in Drosophila,” Aging Cell, 25(2). Available at: 10.1111/acel.70405.

Ikeya, T. et al. (2009) “The endosymbiont *Wolbachia* increases insulin/IGF-like signalling in Drosophila,” Proceedings of the Royal Society B: Biological Sciences, 276(1674), pp. 3799–3807. Available at: 10.1098/rspb.2009.0778.

Kapahi, P. et al. (2004) “Regulation of Lifespan in Drosophila by Modulation of Genes in the TOR Signaling Pathway,” Current Biology, 14(10), pp. 885–890. Available at: 10.1016/j.cub.2004.03.059.

Kapahi, P., Kaeberlein, M. and Hansen, M. (2017) “Dietary restriction and lifespan: Lessons from invertebrate models,” Ageing Research Reviews, 39, pp. 3–14. Available at: 10.1016/j.arr.2016.12.005.

Kuklinski, K.T. et al. (2026) “Innate immune signaling mediates differential acute and long-term outcomes of repeated vs single TBI in Drosophila,” GENETICS, 232(1). Available at: 10.1093/genetics/iyaf249.

Legan, S.K. et al. (2008) “Overexpression of Glucose-6-phosphate Dehydrogenase Extends the Life Span of Drosophila melanogaster,” Journal of Biological Chemistry, 283(47), pp. 32492–32499. Available at: 10.1074/jbc.M805832200.

Ling, D. and Salvaterra, P.M. (2011) “Robust RT-qPCR Data Normalization: Validation and Selection of Internal Reference Genes during Post-Experimental Data Analysis,” PLoS ONE, 6(3), p. e17762. Available at: 10.1371/journal.pone.0017762.

Mathew, R., Pal Bhadra, M. and Bhadra, U. (2017) “Insulin/insulin-like growth factor-1 signalling (IIS) based regulation of lifespan across species,” Biogerontology, 18(1), pp. 35–53. Available at: 10.1007/s10522-016-9670-8.

Mawhinney, R.M.S. and Staveley, B.E. (2011) “Expression of GFP can influence aging and climbing ability in Drosophila,” Genetics and Molecular Research, 10(1), pp. 494–505. Available at: 10.4238/vol10-1gmr1023.

McClure, C.D. et al. (2022) “An auxin-inducible, GAL4-compatible, gene expression system for Drosophila,” eLife, 11. Available at: 10.7554/eLife.67598.

McGuire, S.E. et al. (2003) “Spatiotemporal Rescue of Memory Dysfunction in Drosophila,” Science, 302(5651), pp. 1765–1768. Available at: 10.1126/science.1089035.

Miquel, J. et al. (1976) “Effects of temperature on the life span, vitality and fine structure of Drosophila melanogaster,” Mechanisms of Ageing and Development, 5, pp. 347–370. Available at: 10.1016/0047-6374(76)90034-8.

Mołoń, M. et al. (2020) “Effects of Temperature on Lifespan of Drosophila melanogaster from Different Genetic Backgrounds: Links between Metabolic Rate and Longevity,” Insects, 11(8), p. 470. Available at: 10.3390/insects11080470.

Osterwalder, T. et al. (2001) “A conditional tissue-specific transgene expression system using inducible GAL4,” Proceedings of the National Academy of Sciences, 98(22), pp. 12596–12601. Available at: 10.1073/pnas.221303298.

Partridge, L., Piper, M.D.W. and Mair, W. (2005) “Dietary restriction in Drosophila,” Mechanisms of Ageing and Development, 126(9), pp. 938–950. Available at: 10.1016/j.mad.2005.03.023.

Piper, M.D.W. and Partridge, L. (2016) “Protocols to Study Aging in Drosophila,” pp. 291–302. Available at: 10.1007/978-1-4939-6371-3_18.

Poirier, L. et al. (2008) “Characterization of the Drosophila Gene-Switch system in aging studies: a cautionary tale,” Aging Cell, 7(5), pp. 758–770. Available at: 10.1111/j.1474-9726.2008.00421.x.

Proshkina, E.N. et al. (2015) “Basic mechanisms of longevity: A case study of Drosophila pro-longevity genes,” Ageing Research Reviews, 24, pp. 218–231. Available at: 10.1016/j.arr.2015.08.005.

Regan, J.C. et al. (2016) “Sex difference in pathology of the ageing gut mediates the greater response of female lifespan to dietary restriction,” eLife, 5. Available at: 10.7554/eLife.10956.

Rera, M., Clark, R.I. and Walker, D.W. (2012) “Intestinal barrier dysfunction links metabolic and inflammatory markers of aging to death in Drosophila,” Proceedings of the National Academy of Sciences, 109(52), pp. 21528–21533. Available at: 10.1073/pnas.1215849110.

Richhariya, S. et al. (2023) “Dissecting neuron-specific functions of circadian genes using modified cell-specific CRISPR approaches,” Proceedings of the National Academy of Sciences, 120(29). Available at: 10.1073/pnas.2303779120.

Richhariya, S. et al. (2025) “Metabolic rewiring prevents neurodegeneration caused by chronic mitochondrial dysfunction,” Current Biology, 35(22), pp. 5443–5459.e5. Available at: 10.1016/j.cub.2025.09.063.

Schindelin, J. et al. (2012) “Fiji: an open-source platform for biological-image analysis,” Nature Methods, 9(7), pp. 676–682. Available at: 10.1038/nmeth.2019.

Scialo, F. et al. (2016) “Practical Recommendations for the Use of the GeneSwitch Gal4 System to Knock-Down Genes in Drosophila melanogaster,” PLOS ONE, 11(8), p. e0161817. Available at: 10.1371/journal.pone.0161817.

Simonsen, A. et al. (2008) “Promoting basal levels of autophagy in the nervous system enhances longevity and oxidant resistance in adult Drosophila,” Autophagy, 4(2), pp. 176–184. Available at: 10.4161/auto.5269.

Slack, C. et al. (2011) “dFOXO-independent effects of reduced insulin-like signaling in Drosophila,” Aging Cell, 10(5), pp. 735–748. Available at: 10.1111/j.1474-9726.2011.00707.x.

Sobrido-Cameán, D. et al. (2025) “Mitochondrial ROS and HIF-1α signaling mediate synaptic plasticity in the critical period,” PLOS Biology, 23(8), p. e3003338. Available at: 10.1371/journal.pbio.3003338.

Sun, Y. et al. (2013) “Aging Studies in Drosophila Melanogaster,” pp. 77–93. Available at: 10.1007/978-1-62703-556-9_7.

Tatar, M. et al. (2001) “A Mutant Drosophila Insulin Receptor Homolog That Extends Life-Span and Impairs Neuroendocrine Function,” Science, 292(5514), pp. 107–110. Available at: 10.1126/science.1057987.

Ulgherait, M. et al. (2014) “AMPK Modulates Tissue and Organismal Aging in a Non-Cell-Autonomous Manner,” Cell Reports, 8(6), pp. 1767–1780. Available at: 10.1016/j.celrep.2014.08.006.

Vargas, M.A. et al. (2010) “A Role for S6 Kinase and Serotonin in Postmating Dietary Switch and Balance of Nutrients in D. melanogaster,” Current Biology, 20(11), pp. 1006–1011. Available at: 10.1016/j.cub.2010.04.009.

Wong, R. et al. (2009) “Quantification of Food Intake in Drosophila,” PLoS ONE, 4(6), p. e6063. Available at: 10.1371/journal.pone.0006063.

Yamada, R. et al. (2017) “Mifepristone Reduces Food Palatability and Affects Drosophila Feeding and Lifespan,” The Journals of Gerontology Series A: Biological Sciences and Medical Sciences, 72(2), pp. 173–180. Available at: 10.1093/gerona/glw072.

Zhang, X.-R. et al. (2022) “An improved auxin-inducible degron system for fission yeast,” G3 Genes|Genomes|Genetics, 12(1). Available at: 10.1093/g3journal/jkab393.

Ziehm, M., Piper, M.D. and Thornton, J.M. (2013) “Analysing variation in Drosophila aging across independent experimental studies: a meta-analysis of survival data,” Aging Cell, 12(5), pp. 917–922. Available at: 10.1111/acel.12123.

